# Rethinking mortality rates in men and women: do men age faster?

**DOI:** 10.1101/179846

**Authors:** Peter Lenart, Daniela Kuruczova, Peter K. Joshi, Julie Bienertová-Vašků

## Abstract

Women on average live longer than men, which seems to suggest that women also age slower than men. However, the potential difference in the pace of aging between the sexes is a relatively controversial topic, and both positions, i.e. “men age faster” and “men and women age at the same pace”, have found some support. We therefore employ parametric models previously established in model organisms as well as two nonparametric approaches to compare the pace of aging between the sexes using freely available mortality data from 13 developed countries. Our results support the hypothesis that men age faster than women while also suggesting that the difference is small and that from a practical standpoint male mortality rates behave similarly to the mortality rates of women approximately eight years their senior.

## Introduction

The life expectancies of men and women are widely recognized as being different: women worldwide live longer than men. This logically leads to the question whether women also age slower than men. Both “yes” and “no” answers have found some support.^1–3^ The classical argument against the notion that women age slower is the fact that men experience higher mortality rates at almost every age, i.e. that the reason for their shorter lifespan is that men are the less “robust” sex and as such exhibit higher background mortality.^1,3^ On the other hand, researchers suggesting that women age slower than men note that this line of reasoning may not be altogether valid since men die from different causes at different ages.^2^ Regardless of theoretical arguments, aging can be defined as an age-dependent increase in mortality ^2,4,5^ and the pace of aging of men and women may therefore be empirically calculated using available mortality data. In this article, we present the results of a mortality data analysis which suggests that men do age faster than women – once a sex-specific baseline mortality level is accounted for.

One method of quantifying aging relies on calculating the rate at which mortality increases with age.^6^ The relationship between human age and mortality is usually modeled using a predefined distribution which explicitly defines the relationship between age and mortality rate. Distributions most commonly used for this purpose include Gompertz, its extension Gompertz– Makeham, Weibull or logistic.^7^ The choice of a specific distribution depends on the purpose of its use: the best-fitting model is often desired when a prediction is sought while a different model may be more suitable for the interpretation of parameter values.^8,9^ Since our objective was to test the difference between the pace of mortality rate increase in men and women, the Gompertz model^10^ was selected as a simple and suitable option. In addition to accommodating human mortality data between approximately 30 and 80 years of age,^9,11^ it also offers a means for comparing mortality rate increase by means of mortality rate doubling time (MRDT), a parameter commonly used as an estimate of the rate of aging.^12,13^ On the other hand, it does not distinguish between intrinsic and extrinsic mortality rates, where intrinsic mortality is assumed to be the result of aging and increases over time while extrinsic mortality is assumed to be caused by environmental hazards and is thus constant over time.^14^ However, this inability to distinguish between intrinsic and extrinsic mortality rates is to some extent alleviated by the fact that mortality within the chosen interval of 30 to 60 years of age is mainly influenced by intrinsic causes.^9^ The Gompertz–Makeham model extends the Gompertz model to include mortality rate independent of age. This partitioning of mortality rates into an age-related and a constant component is clearly helpful when analyzing the rates of aging.

In this study we used mortality data obtained from the Human Mortality Database^15^ to calculate MRDTs using the Gompertz and Gompertz–Makeham model for male and female populations in 13 developed countries. Furthermore, we have also employed two non-parametric approaches to compare the pace of aging between sexes. However, it must be said that mortality rates are affected by a great variety of external influences unrelated to aging. One extreme example of such external influences was undoubtedly World War II, which dramatically altered mortality rates both directly through the deaths of millions of soldiers and civilians and indirectly through the late effects of injuries, starvation, psychological trauma, etc. It is known that mortality rates during the early life of a cohort influence its mortality rates later in life^16^ which makes cohorts affected by a WWII unsuitable for comparing the pace of aging between sexes. Because most countries in the Human Mortality Database were more or less heavily involved in WWII, we analyzed mortality patterns only in people born at least five years after the end of this conflict. To be more specific, we analysed cohorts of people born from 1950 to 1954 using their mortality rates in periods starting from 1980 to 1984 to the newest available data in the Human Mortality Database. In other words, investigated mortality rates were calculated using periods starting with the subjects’ 30^th^ birthdays and ending with the end of records.

## Methods

Mortality rate data were acquired from www.mortality.org on 12 July 2017. The Human Mortality Database (HMD) contained data about mortality rates for 39 sovereign countries and several others smaller areas and populations. In our analysis we focused on 13 developed, western (plus Japan), stable countries with populations exceeding 8 million. The analysed countries are: Australia, Belgium, Canada, France, Italy, Japan, the Netherlands, Portugal, Sweden, Switzerland, the United Kingdom, the United States of America and West Germany.

### Gompertz and Gompertz-Makeham model

The Gompertz model^10,12^ of exponential hazard growth was used to model the relationship between age and mortality rate. The basic form of the Gompertz model is

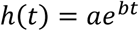

where *a* and *b* are constants, *t* is time (age), and *h(t)* is the hazard (mortality) rate. Using the logarithmic transformation, a simple linear model is obtained

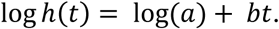

where *log(a)* signifies the intercept (overall shift of the line in the direction of the y-axis) and *b* expresses the slope of the line. MRDT is subsequently calculated from the slope as

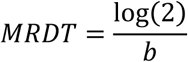

and expresses the time it takes for the mortality rate to double.

The Gompertz–Makeham model is a natural extension of the Gompertz model obtained by adding a constant:^17^

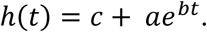

The constant *c* expresses the part of mortality that does not depend on age. Focusing only on the age-dependent part of the equation, the mortality rate doubling time can be obtained in the same manner as in the Gompertz model, using the value of parameter *b*.

We fitted the above described on data for each individual country using an age interval beginning at 30 years of age. We fitted one model for both male and female data with sex as a dummy variable in order to compare them. Due to the exponential nature of models, we used numerical fitting using non-linear least squares. Both models accurately fit human mortality dynamics roughly between 30 and 80,^9,11^ which we subsequently confirmed using an exploratory analysis of HMD data.

### Nonparametric approaches

#### Smoothing spline approach

We also used a nonparametric approach to model the relationship between age and mortality rate. We approximated the death rate data by a smoothing spline^18^ (smoothing parameter 0.7) and subsequently calculated the derivative curve. The value of the derivative at each time point expresses the rate of mortality increase. The derivative value would be zero for a constant mortality rate, positive values indicate an increase in the mortality rate over time – the higher the derivative value, the steeper the mortality curve.

#### Mortality rate matching approach

To gain more intuitive insight into the difference between male and female mortality rates we matched the female mortality rates with male mortality rates. To ensure the continuity of both male and female mortality rates over time, we used mortality rate curves obtained using the previous approach. The resulting line expresses the ages where the female mortality rate equals the male mortality rate.

## Results

### Gompertz model

MRDTs calculated for people born in 1954 are longer for males in 10 out of 13 countries (Figure S1 and Table S1). However, the possibility of longer male MRDTs is inconsistent with MRDTs calculated for 1953, 1952, 1951 and 1950 cohorts. Males born in 1953 have longer MRDTs in 7 out of 13 countries but those born in 1952 only in 6 out of 13. Furthermore, males born in 1951 and 1950 have longer MRDTs only in 8 and 7 countries respectively. Gompertz model results thus suggest that MRDTs are the same for males and females.

### Gompertz–Makeham model

Contrary to the results of the Gompertz model, MRDTs calculated using the Gompertz– Makeham model for 1950–1954 cohorts exhibit consistent differences between the sexes (Figure 1 and Table S2). MRDTs for the 1954 cohort are longer for women in 11 out of 13 countries while MRDTs for the 1953 cohort are longer for women in 12 countries. This trend is further evident in all remaining cohorts. MRTDs for the 1952, 1951 and 1950 cohorts are higher for females in 11, 10 and 11 out of 13 countries respectively.

**Figure 1:**
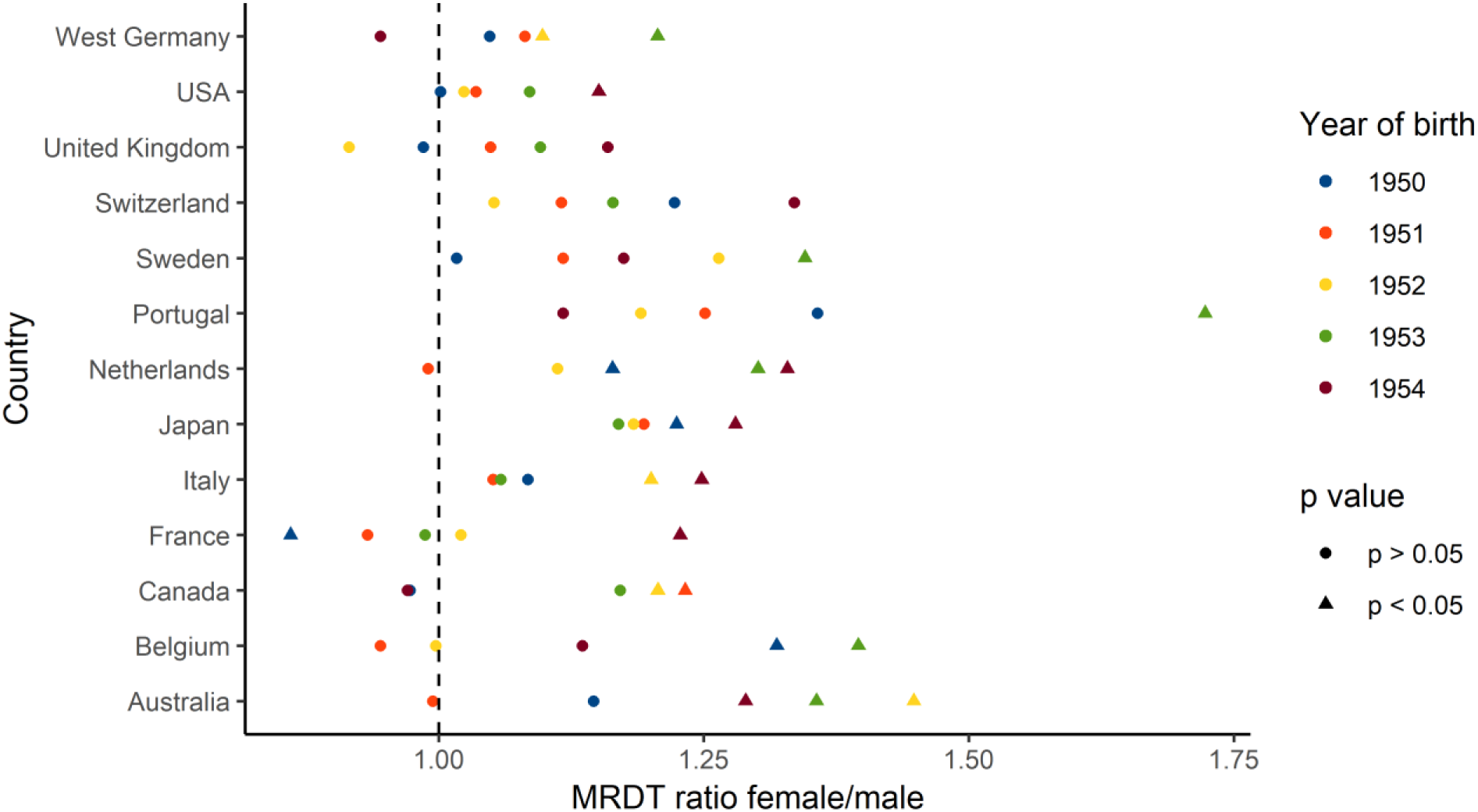
Mortality rate increase is greater for men than women. The figure provides a summary of the ratio between female and male MRTDs calculated using the Gompertz–Makeham model for all analyzed countries and birth cohorts.

The disparate outcomes of the Gompertz and Gompertz–Makeham models are most likely caused by different parametrization rather than by different curve shapes (Fig. 2). When the age-independent parameter c is not included, like in the Gompertz model, the value of the other two parameters changes accordingly in order to provide a best-fitting curve. If a roughly similar mortality curve were to be described by the Gompertz and Gompertz–Makeham models, the following would apply: the growing c value of the Gompertz–Makeham corresponds to a decreasing b value in the Gompertz model and thus an increasing MRDT. We have found that the age-independent parameter c is higher in males in most cases (Table S2) which partially explains the different results achieved using the Gompertz and Gompertz–Makeham models.

**Figure 2:**
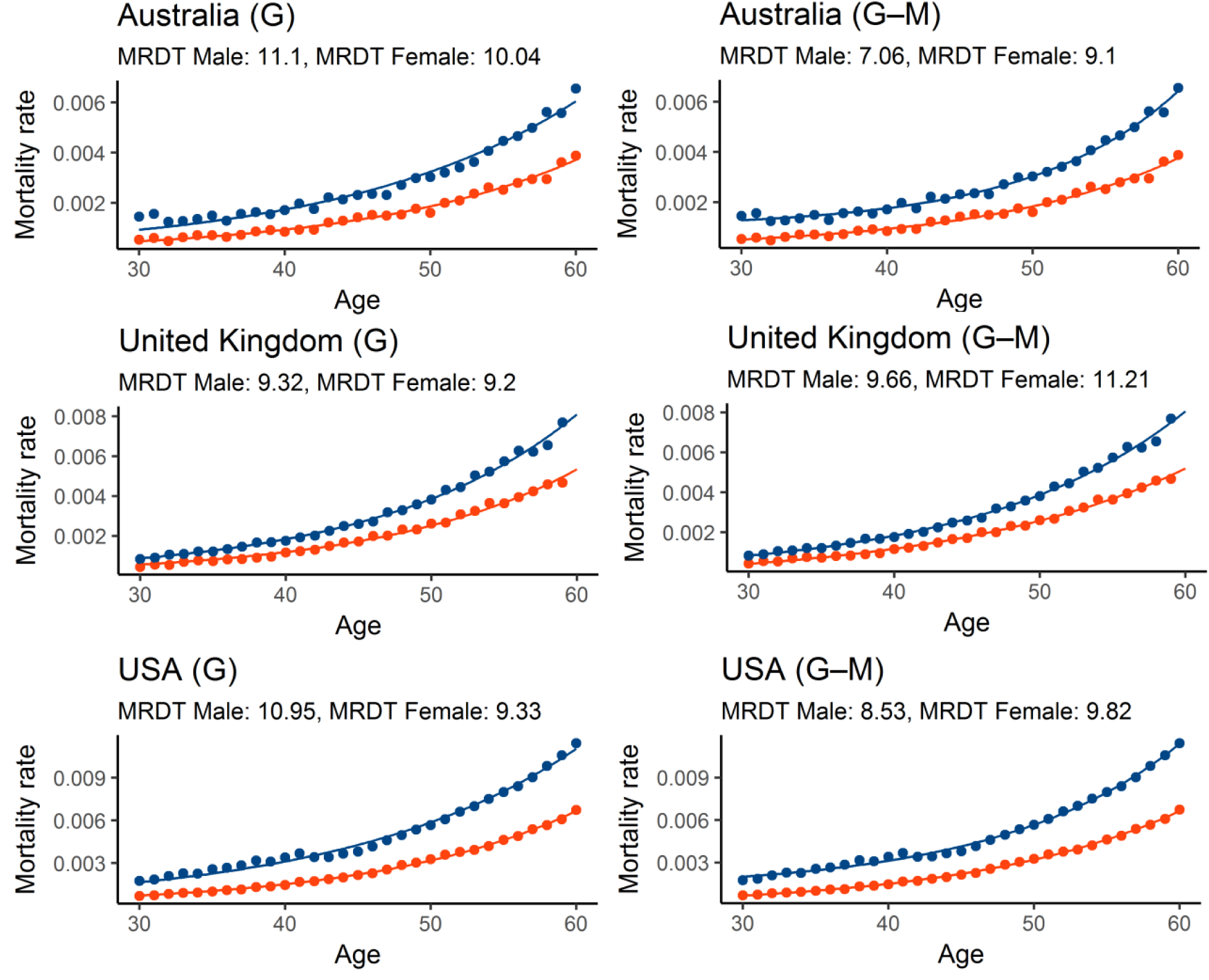
Gompertz and Gompertz–Makeham models provide similar fit. **A** comparison between curve shapes for Gompertz (G) and Gompertz Makeham (G–M) in three selected countries. Blue represents male and red female mortality curves.

### Smoothing spline approach

We employed a nonparametric approach to further test the difference in the pace of aging between the sexes without the constraint of parametric models. Results in the form of derivative curves, where the value of the derivative at each time point constitutes an increase of mortality rate per year, (Figure 3 and Table S3) clearly show that mortality rates increase faster in men than in women in all studied countries and cohorts. These results are thus in agreement with the results of the Gompertz–Makeham model.

**Figure 3:**
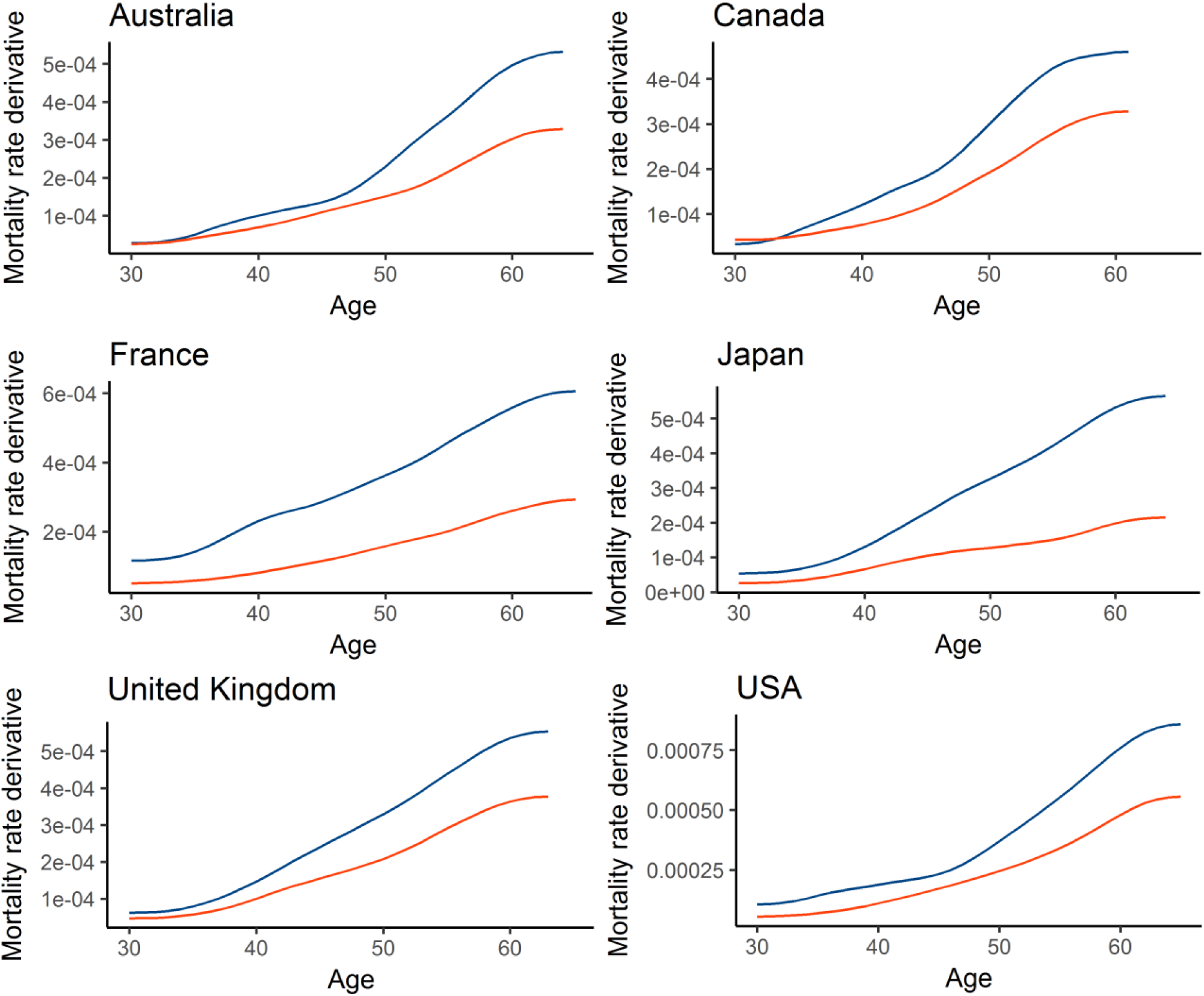
Mortality curves of men are steeper than those of women. The figure shows derivative curves obtained for men and women using a nonparametric approach for six selected countries in the 1950 cohort. Blue mortality curves are fore men while red for female.

Furthermore, a comparison of derivative curves for males and females enables us to quantify the ratio of annual male and female mortality rate increase (Figure 4). The median derivative ratio for males and females ranges from 1.34 in the Netherlands to 2.36 in Japan (Figure 5). In other words, the median increase in male mortality rates for this cohort is 34 to 136 % higher than that in females. The median derivative ratio for the 1951 cohort ranges from 1.37 for the Netherlands to 2.56 for Portugal. The ranges for 1952, 1953 and 1954 cohorts are 1.28–2.68, 1.33–2.87, and 1.3–2.38 respectively.

**Figure 4:**
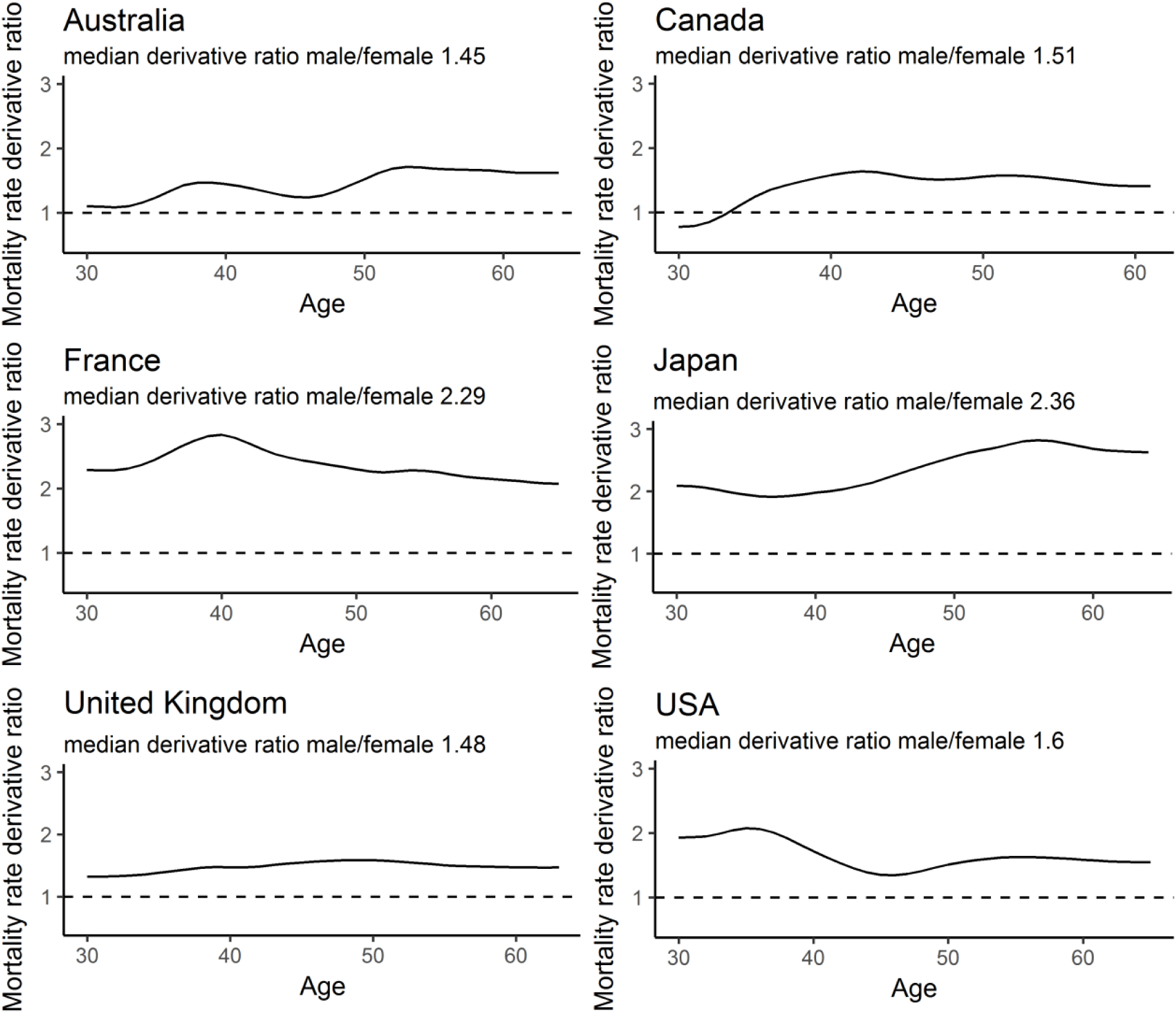
Quantitative differences in male and female mortality rate increase. The figure shows male/female death rate derivative ratios for 1950 in six selected countries.

**Figure 5:**
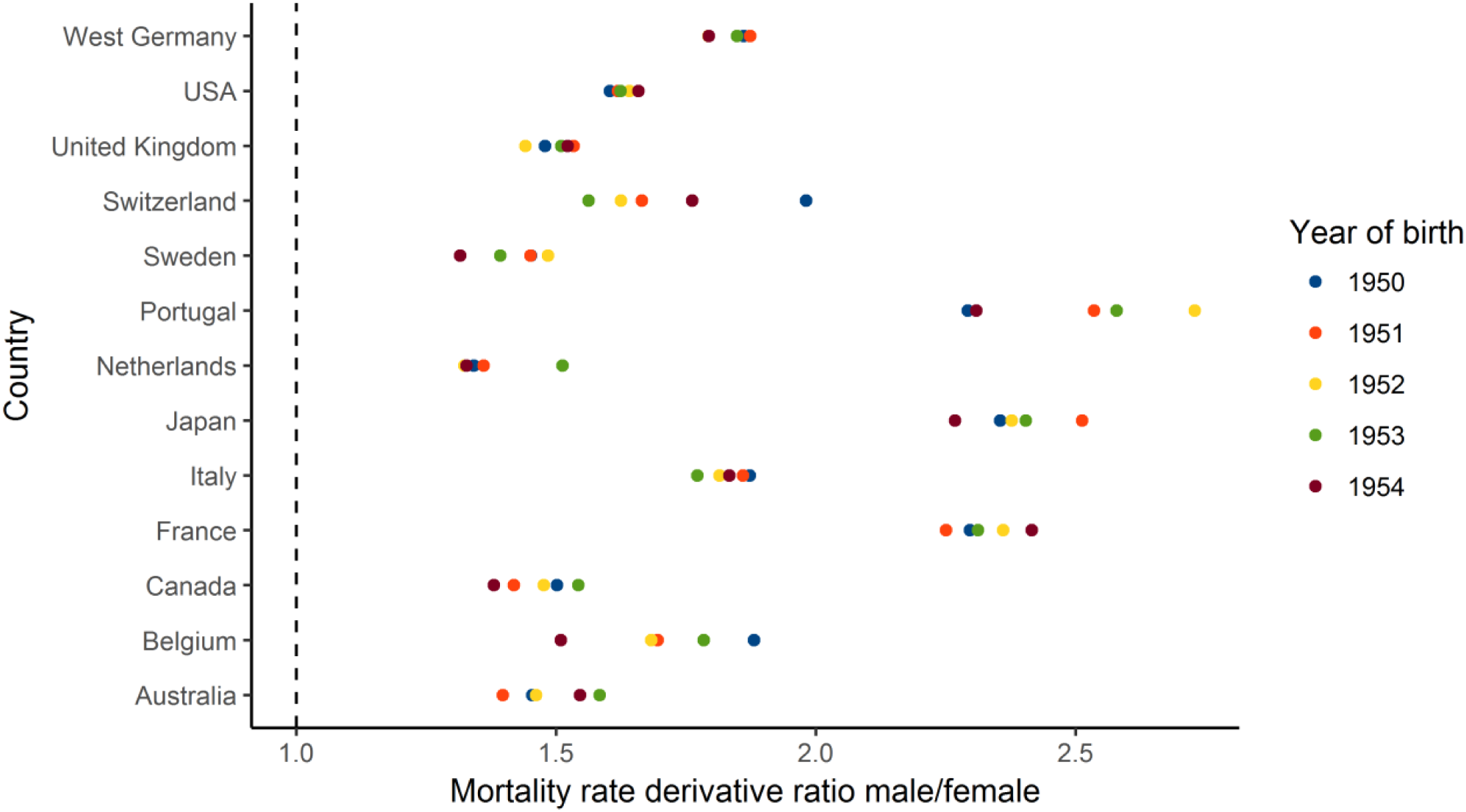
Nonparametric approach shows universally higher increase in male mortality. The figure shows male/female derivative rations for all studied countries and cohorts.

### Mortality rate matching approach

Since the selected calculation methods may have influenced our results, we have decided to employ a second nonparametric approach. By comparing absolute mortality rate values rather than derivatives, we tested whether our results are robust enough to withstand various methodological approaches. When comparing the mortality rates of males and females, we found that the mortality rates of females aged 50 are on average equal to the mortality rates of males aged 41.9 years (Figure 6 and Table S4). When we made the same comparison for the mortality rates of women aged 60 we found out that men had already achieved equal mortality rates at the average age of 51.2. In other words, the gap between female and male mortality rates increased on average by 0.7 years over the course of the decade (Figure 6 and Table S4), suggesting that males age faster.

**Figure 6:**
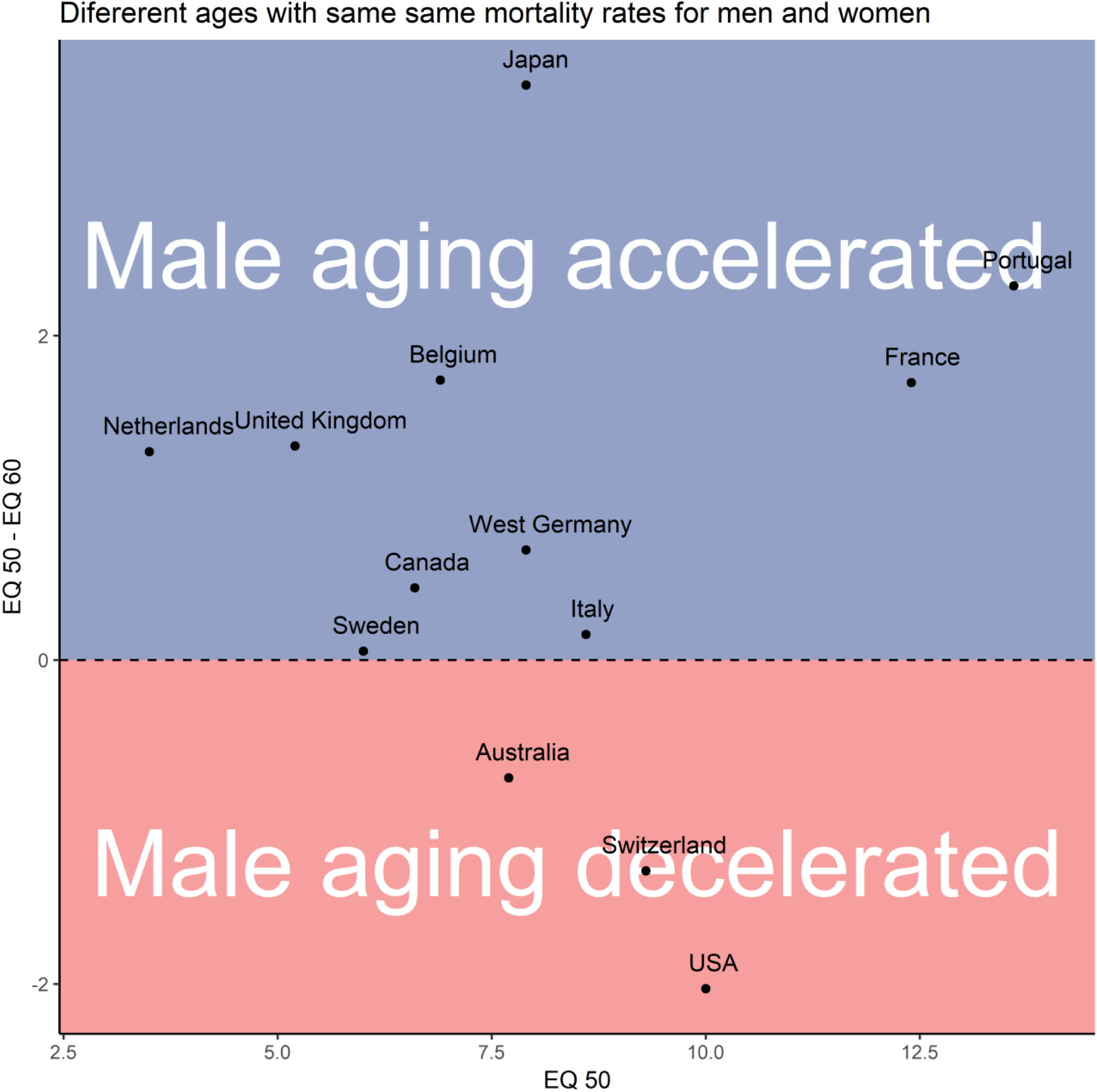
Men achieve mortality rates equal to that of women aged 50 many years earlier. Equivalent mortality (EQ) at age X expresses how much earlier men achieve the same mortality rates as women aged X. For example, an EQ 50 of 10 means that men aged 40 exhibit the same mortality rates as women aged 50. The y axis captures how differences in age with the same mortality rates developed over a ten-year-long period in the studied countries and cohorts.

On the other hand, when we compared the age difference between equal male and female mortality rates using longer timescales, it seems that the observed changes are rather small and that male mortality rates have a dynamic similar to the mortality rates of females several years older (Figure 7).

**Figure 7:**
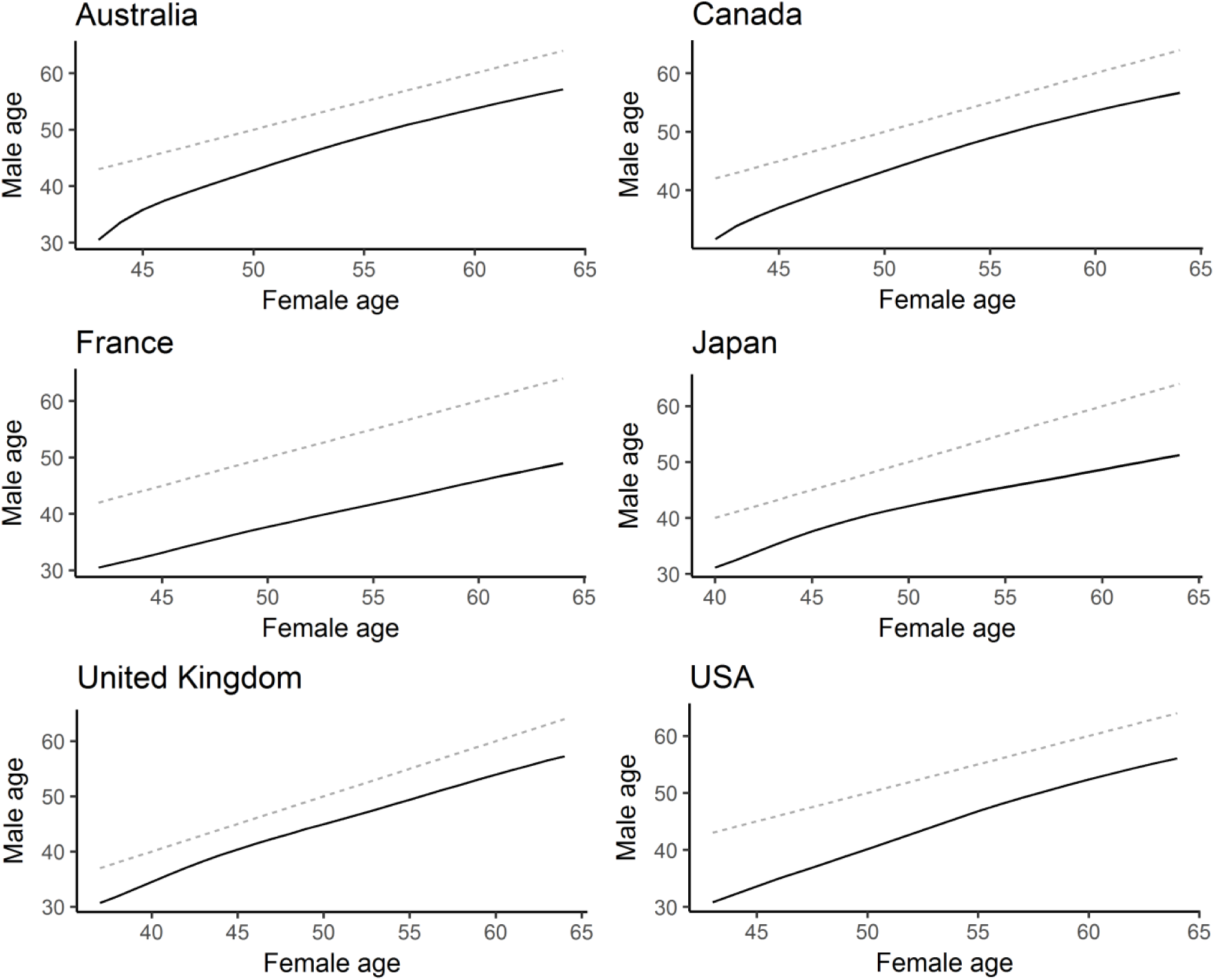
Male mortality rates behave like mortality rates of females several years older. The bottom line in each graph connects ages at which males and females exhibit identical mortality rates. The dotted lines represent a hypothetical situation where males exhibit the same mortality rates as females of the same age.

## Discussion

We investigated mortality data to test whether men age faster than women, as previously suggested in several studies.^2,19–21^ While calculations MRDTs conducted using the Gompertz model do not show a consistent difference in the pace of aging between the sexes, the results of the Gompertz–Makeham model strongly suggests that men age faster than women. The difference between the results of the Gompertz and Gompertz–Makeham model may be partially explained by the fact that, unlike the Gompertz model, the Gompertz–Makeham model includes an age-independent parameter c which is generally higher in men. Thus, in Gompertz model age-independent mortality attributes to MRDTs and inflates male MRDTs more than female MRDTs, which implies that the Gompertz–Makeham model is better suited to comparing aging between the sexes. Nevertheless, although the Gompertz and Gompertz–Makeham models are certainly useful, they are ultimately highly constrained models which may sometimes produce incorrect results. We therefore also compared the pace of aging between the sexes using non-parametric approaches. The results of the smoothing spline approach suggest that men age faster than women, i.e. they are in agreement with with the outcome of the Gompertz–Makeham model. On the other hand, the results of the mortality matching rates approach show that while men may age faster on average, the difference between the male and female paces of aging is rather small. Furthermore, the results of the mortality matching rates approach show that male mortality rates behave similarly to the mortality rates of females approximately eight years older.

While our analysis of mortality data does not distinguish between intrinsic and extrinsic sources of mortality, this partitioning has been examined in several existing studies^22,23^. Despite the fact that mortality partitioning remains a gold standard which may help bring important insight into the aging process in many situations, it is in fact rather superficial: the assumption that intrinsic mortality sources are caused by aging while extrinsic mortality sources are caused by environmental influences – and are thus constant over time – is simply wrong.^22^ Accordingly, even Bruce A. Carnes and S. Jay Olshansky, arguably the two most influential authors studying mortality partitions, sharply disagree with this naive assumption. This is probably best documented by the fact that both are among the authors of a paper which clearly states that “It is difficult to envision a cause of death for humans or any other species, either intrinsic or extrinsic, that does not exhibit age-dependence.”^22^ Furthermore, it is documented that even mortality caused by accidents such as falls, drowning, transport accidents and exposure to mechanical forces dramatically increases with age, as does the number of deaths caused by natural disasters, including excessive heat or cold, earthquakes, lightning, storms, and floods.^14^ In other words, biologically older individuals are at a higher risk of death from both intrinsic and extrinsic sources. We thus believe that using overall i.e. non-partitioned mortality to compare the pace of aging should be sufficient or even preferable to focusing purely on intrinsic mortality.

Though not entirely conclusive, our results support the hypothesis that men age faster than women. This may be important since the faster aging of men could have far-reaching implications for aging research as well as for medicine. Furthermore, while the survival advantage of females has been reported in other primates, it is not typical of all mammals.^3^ If we therefore assume that the longer lifespan of one sex should to some extent reflect differences in aging between the sexes, it also implies that the faster aging of men is a recent evolutionary development. In case this interpretation is correct, it then leads to an obvious question: what triggered the evolution of sex-specific aging rates?

On the other hand, our study demonstrates that a comparison of the paces of aging may yield vastly different results when different methods are employed (e.g., Gompertz vs. Gompertz–Makeham), which is an issue deserving of broader scientific attention. Furthermore, our results show that if men and women age at different paces, the difference is rather small and it seems that, from a practical viewpoint, male mortality rates behave in the same way as the mortality rates of women several years older.

## Code availability

Code is available at http://www.math.muni.cz/∼xkuruczovad/Mortality/ or upon request.

## Supporting information

## Acknowledgments

The project was supported by the CETOCOEN PLUS (CZ.02.1.01/0.0/0.0/15_003/0000469) project of the Ministry of Education, Youth and Sports of the Czech Republic. The project was also supported by the RECETOX Research Infrastructure (LM2015051 and CZ.02.1.01/0.0/0.0/16_013/0001761). Furthermore, Peter Lenart received support from the Brno Ph.D. Talent competition.

## Authors’ contributions

P. L. formulated the research problem, chose data sources and interpreted the results. D. K. analyzed the data P.K.J, and J. B. V supervised the project. All authors co-wrote the manuscript.

## Competing interests

The authors declare no competing interests.

